# Molecular phylogeny of wood decay fungi of hardwood and their ability to produce laccase that correlates with triphenylmethane dye decolorization

**DOI:** 10.1101/648147

**Authors:** Maduri Piumi Sashikala Mahawaththage Dona, Anushi Suwanethya Deraniyagala, Priyanga Wijesinghe, Renuka Nilmini Attanayake

## Abstract

Though Sri Lanka belongs to one of the 34 biodiversity hotspots of the world, its microfolora specially fungi are not well studied and underrepresented in the global literature. Here we report the fungal species diversity of decaying hardwood of a Sri Lankan dry zone forest for the first time. Decaying hardwoods were collected from historically important Dimbulagala forest reserve, Sri Lanka and fungi associated with these woods were isolated. Out of 35 fungal species identified using morphological and molecular methods, 11 species were first records in Sri Lanka. All the tested isolates were able to utilize wood as the sole carbon source and produced varying degrees of laccase. Isolates of *Perenniporia tephropora, Coriolopsis caperata, Gymnopilus dilepis, Fusarium solani* and *Vanderbylia fraxinea* were among the top six laccase producers. Except *Fusarium solani*, the rest of the isolates showed more than 70% decolorization of the of triphenylmethane dye and there was a significant positive correlation between laccase production and dye decolorization. To the best of our knowledge laccase production and dye decolorization ability of *Vanderbylia fraxinea* and *Gymnopilus dilepis* have never been reported in the fungal kingdom before. *Perenniporia tephropora* was isolated from one of the strongest decay resistant hardwood species, Ebony (*Diospyros ebenum*) also known as dark wood and *V. fraxinea* was isolated from another medicinally important hardwood Neem (*Azadirachta indica*). Findings of this study confirms that decaying hardwood of Sri Lanka provide unexplode a unique niche for discovering fungal species with biotechnological applications such as high laccase producers and dye decolorizers.

## Introduction

Fungi are considered to be miniature metabolic factories having versatile secondary metabolites and enzymes with the potential to be used as work horses in diverse array of biotechnological applications [1]. Based on the Hawksworth’s [2] estimation, there are abut of 1.5 million fungal species present on earth, which is considered as the baseline estimation. However, recent metagenomics studies suggested that actual numbers might be closer to 3.5 to 5.1 million species or much higher than this [3]. This uncertainty in the numbers is partially due to lack of advanced molecular based thorough studies in the tropics where incredibly rich diversity has been reported [4]. Hawksworth [5] also suggested that much of the undescribed fungal species could be present in the tropics and it is reviewed in Aime and Brearley [4].

Sri Lanka, a tropical island in the Indian ocean along with the Western Ghats, belongs to one of the 34 biodiversity hotspots in the world. Though its plant and animal diversity is well studied [6,7], microbial studies, especially fungal studies remain in its infancy. Furthermore, biodiversity hotspot concept of Sri Lanka should not be an exception for microbial diversity including fungi. However, most of the fungal studies in Sri Lanka have mainly focused on macro-fungi using morphological characters [8]. Moreover, these studies have mainly concentrated on the wet zone forests. On the other hand, dry zone forest ecosystems in the country spread over 22 % of area in Sri Lanka, whereas the total forest cover is about 26.6%. Dimbulagala (7°51’40.5”N 81°07’05.5”E) is an isolated hill covered with dry zone forest and it is rich with strong and economically important hardwood bearing plant species. This region is contacted with minimum anthropological activities for many years mainly due to 30-year long civil war in this region and was the study site of the current study. An unexplored niche, decaying hardwood, of Dimbulagala forest reserve was selected with the aim of describing the species diversity of hardwood decay fungi for the first time in Sri Lanka.

For a healthy forest ecosystem, it is critical to recycle carbon stored in these hardwood and litter. Among many organisms such as beetles, flies, slime molds, bacteria, slugs and snails, primary decomposers that recycle carbon are fungi. It is known that wood, specially hardwood is a challenging substrate to degrade as it has very low nitrogen content (commonly C: N is about 500:1) and the presence of various fungal toxic compounds. Therefore, fungi specially basidiomycetes have evolved to degrade the hardwood structure consisted of cellulose, hemicellulose and the strongest natural polymer, lignin. They secrete major lignin modifying enzyme families such as lignin peroxidase, manganese peroxidase, versatile peroxidase and laccase [9]. Among them, fungal laccases have attracted the attention of scientific community due to its high redox potential and versatile catalytic properties compared to the laccases from other sources such as plants and bacteria [10]. Sri Lankan dry zone forests, including Dimbulagala forest reserve are rich in hardwood bearing plant species such as *Diospyros ebenum, Manilkara hexondra, Vitex pinnata, Diospyros chaetocarpa, Terminalia bellirica, Chloroxylon swietenia* and *Berrya cordifolia* [11–13]. These plant species produce hardwood with high economic value and are resistant to degradation. However, mycelial mats and mushrooms were often observed in decaying hardwoods in these areas indicating the involvement of fungal for the decay process. If that is the case, fungal species found in association with these decaying hardwood will be able to utilize wood as the sole carbon source. Therefore, the second objective was to assess the ability of the fungal isolates to grow on wood dust media. It was hypothesized that fungal species capable of growing on such hardwood have the ability to produce high amounts of laccases.

Fungal laccases have been identified not only as cell wall degraders, but also as strong oxidizers with the ability to oxidize di- and polyphenols, aromatic amines, and detoxify environmental effluents of food, paper, pulp and textile dyes [14,15]. Among these abilities of laccase, industrial dye decolorization has been the subject of many studies [16,17] and laccases are often considered as a “Green Tool” in biotechnology [18]. Basidiomycetes have often found to be high laccase producers as well as dye decolorizers. It was also intended to determine whether the isolates of the current study are capable of decolorizing an industrial dye, triphenylmethane with the hope of identifying novel dye decolorizers.

Out of tested 43 fungal isolates that could utilize wood as the sole carbon source, 11 were first records of Sri Lanka. Out of them two species were first time records as laccase producers and tryphenyl methane dye decolorizers. There was a strong positive correlation between laccase production and dye decolorization.

## Materials and Methods

### Sample collection

Sampling site was Dimbulagala historical forest reserve in Polonnaruwa district, Sri Lanka. Research permits were obtained from the authorities and samples were collected and handled ethically without harming living trees. Thirty-five decayed hard wood pieces of 5-6 cm length showing brown and white rot symptoms were collected randomly along the trail to the top of the hill (7°51’34.1”N 81°06’53.2”E to 7°51’42.8”N 81°07’09.0”E). Samples were collected into clean zip lock plastic bags and transported to the laboratory at University of Kelaniya (Fig. 1). Host of the decaying wood was recorded whenever possible. However, in most of the cases it was not possible to clearly identify the plant species that decayed wood pieces belonged to. Samples were air dried for two days and stored in a refrigerator until further use.

**Fig 1:**
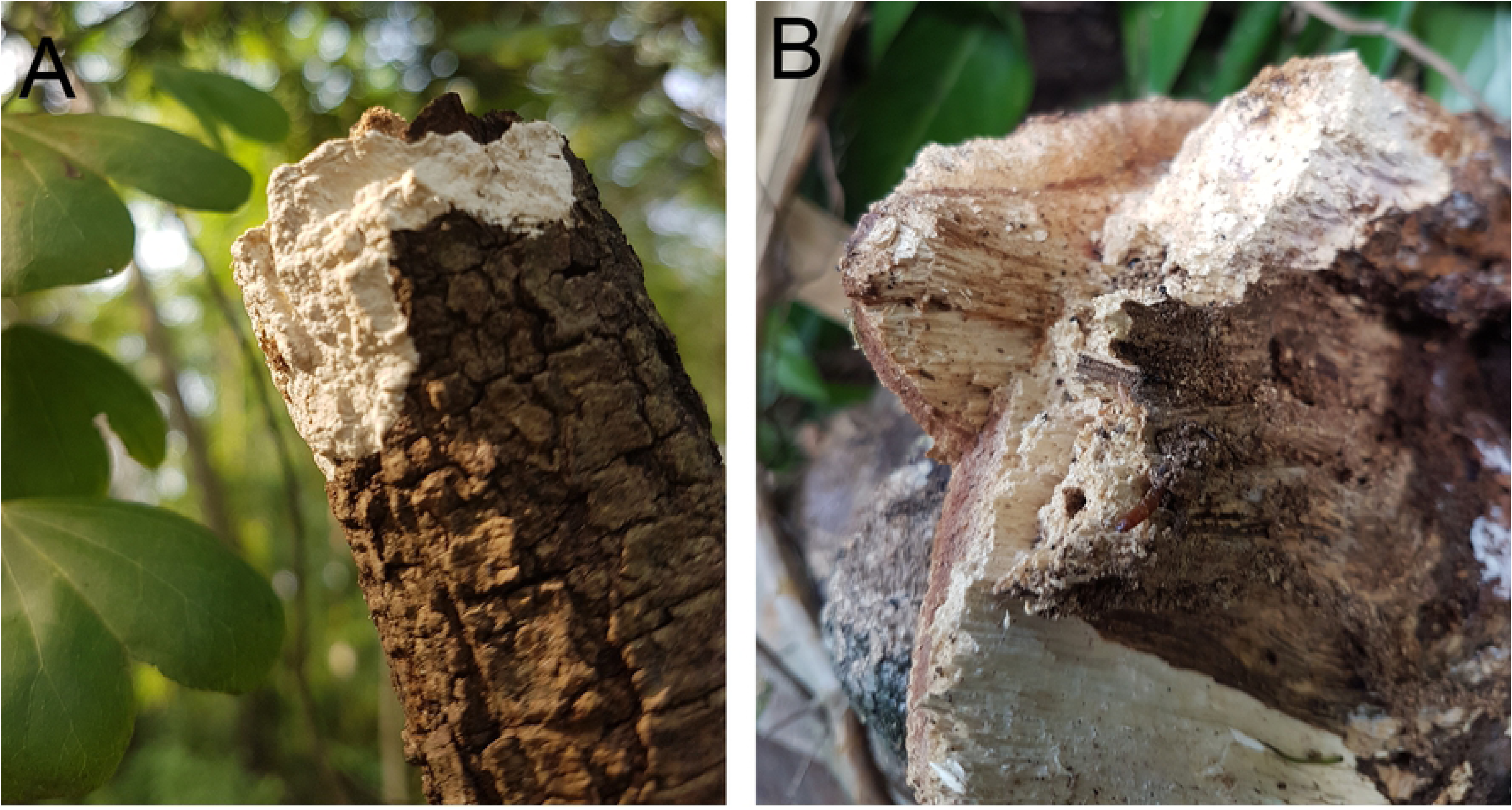
Samples showing decaying hardwood (A) with mycelial growth and (B) white rot symptom.

### Isolation of fungi into culture media

Decaying wood pieces (approximately 1 × 1 cm) were surface sterilized with 5% Clorox for 2 minutes followed by three serial washings with sterilized distilled water. Washed materials were blot dried using sterilized filter papers. Samples were trimmed from the edges and cut into approximately four pieces, placed on semi selective PDA media supplemented with streptomycin (100 µg/mL). Another isolation step was carried on semi selective medium slightly modified from Baumgartener et al. [19] by supplementing PDA with streptomycin (100 µg/mL) and fungicide carbendazim (4 µg/mL) to inhibit fast growing ascomycetes. Plates were incubated at room temperature for one week. All the fungal colonies originated from the wood pieces were sub cultured and pure cultures were obtained by hyphal tip isolation method. Pure cultures of 55 fungal isolates were obtained and vouchered both in sterile water and at −20 °C in Whatman number 1 filter papers at the Department of Botany, University of Kelaniya, Sri Lanka and assigned strain IDs.

### Ability of isolated fungi to utilize wood as the sole carbon source

Ability of isolated fungi to utilize wood as the sole carbon (C) source was tested according to Swe [20]. During this assay most of the isolates those with morphological characters similar to *Trichoderma* spp. were excluded and therefore, out of 55, 43 isolates were used in further studies. First, saw dust of *Mangifera indica* was passed through 2 mm sieve. Wood agar plates were prepared with saw dust (2% w/v), stock salt solution (5 % v/v), Hutner’s trace elements (0.02 % v/v), with and without supplementing glucose (0.05% w/v) and agar (1.5 % w/v) followed by Swe [20]. Both wood agar plates with and without glucose were inoculated with 5 mm-diameter mycelial disc obtained from actively growing edge of 7-day old culture from each isolate. Plates were incubated at the room temperature (28 ± 2 °C) for three days. Colony diameters were measured in two-dimension perpendicular to each other. Experiment was repeated once.

### Morphological and molecular identification of fungi

All the isolates were grown on PDA plates and used for morphological observation. Colony color, appearance and mycelial color, spores and sporulating structures when available of all the isolates were observed under phase contrast microscope (x 400 magnification) (Olympus CX41 model, Tokyo, Japan).

Out of all the isolates, 35 isolates that showed variable morphological features indicating putatively different species were selected for molecular level identification. Three mycelial plugs from actively growing colonies were inoculated into 50 mL Potato Dextrose Broth (PDB) supplemented with streptomycin (100 μg/mL). Flasks were incubated in a rotary shaker (Stuart-SSL1, UK) at 120 rpm at room temperature for 5-7 days. Mycelia were separated, washed with 500 μL of sterilized distilled water and squeeze dried with sterilized filter papers. DNA was extracted following modified method of Cenis [21]. DNA Pellet was re-suspended in TE buffer (50-100 µL) and stored at −20°C until further analysis. Quality of extracted DNA was visually observed on 0.8% agarose containing 0.5 μg/ml of Ethidium bromide. Polymerase Chain Reaction (PCR) was performed with universal ITS1 and 4 or 4 and 5 primer pairs [22]. Reaction mixture consisted of 1x Colorless GoTaq® Flexi Buffer, 2 mM MgCl_2_, 200 µM each dNTP, 0.5 μM forward and reverse primers, 1.25U of GoTaq® DNA Polymerase (Promega Inc., USA) and <0.5 μg template DNA. PCR reaction was performed in a thermal cycler (Veriti® 96-Well Thermal Cycler, ABI Biosystems, USA) as published in Maduranga et al.[23]. Nuclease free water was added as the negative control and PCR products were separated using agarose gel electrophoresis and visualized under a Gel Documentation System (Quantum ST5, Germany). Pure PCR products were sequenced following Sanger dideoxy chain termination technology at the Genetech Institute, Colombo, Sri Lanka. Sequences were manually edited using BioEdit sequence Alignment Editor (Version7.2.5)[24]. Using Basic Local Alignment Search Tool (BLASTn), nucleotide sequences were compared with the authentic sequences available in the National Center for Biotechnology Information (NCBI) database and species were identified. All sequences were deposited in the GenBank and the accession numbers obtained are shown in Table 1.

**Table 1:**
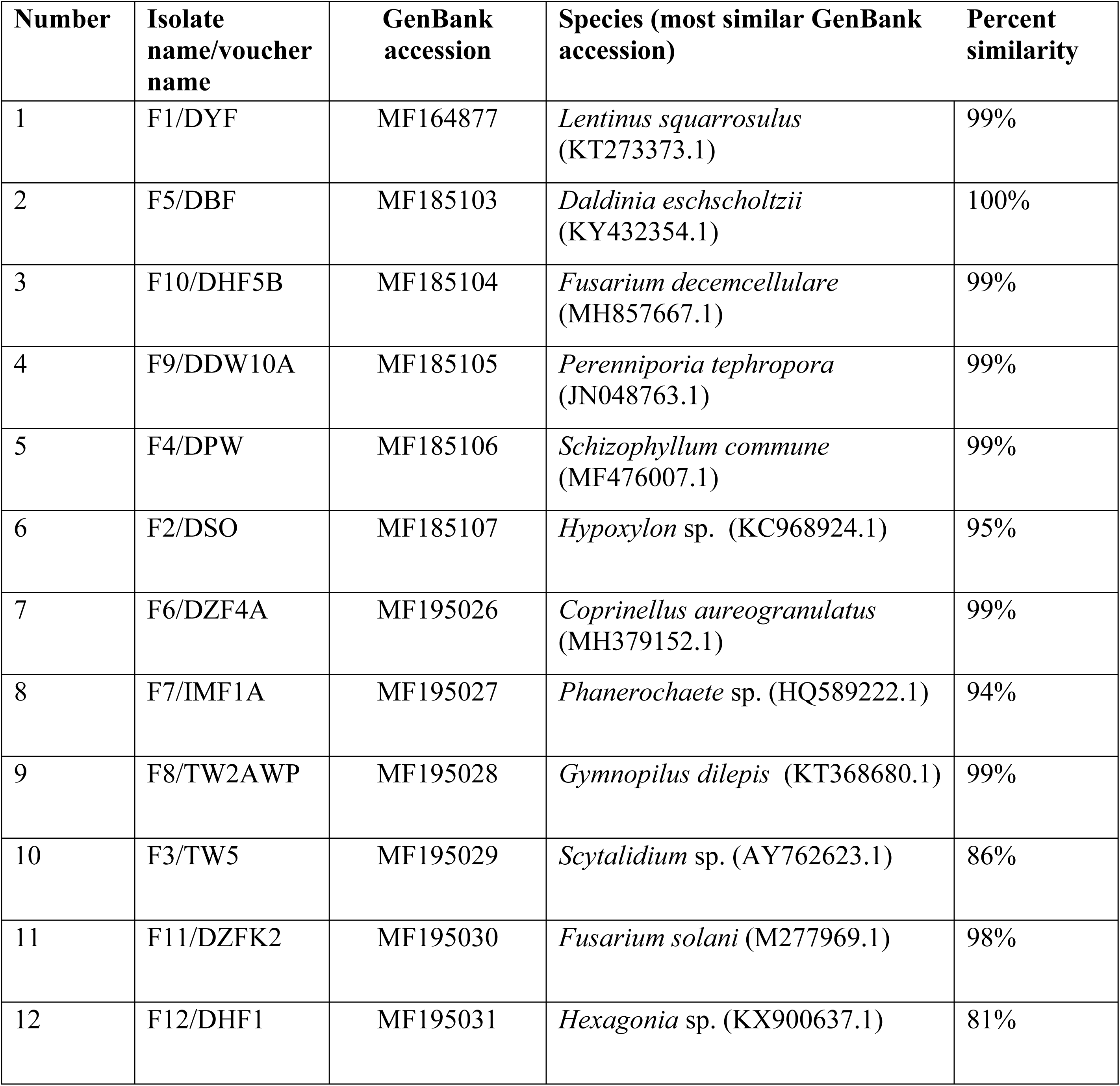
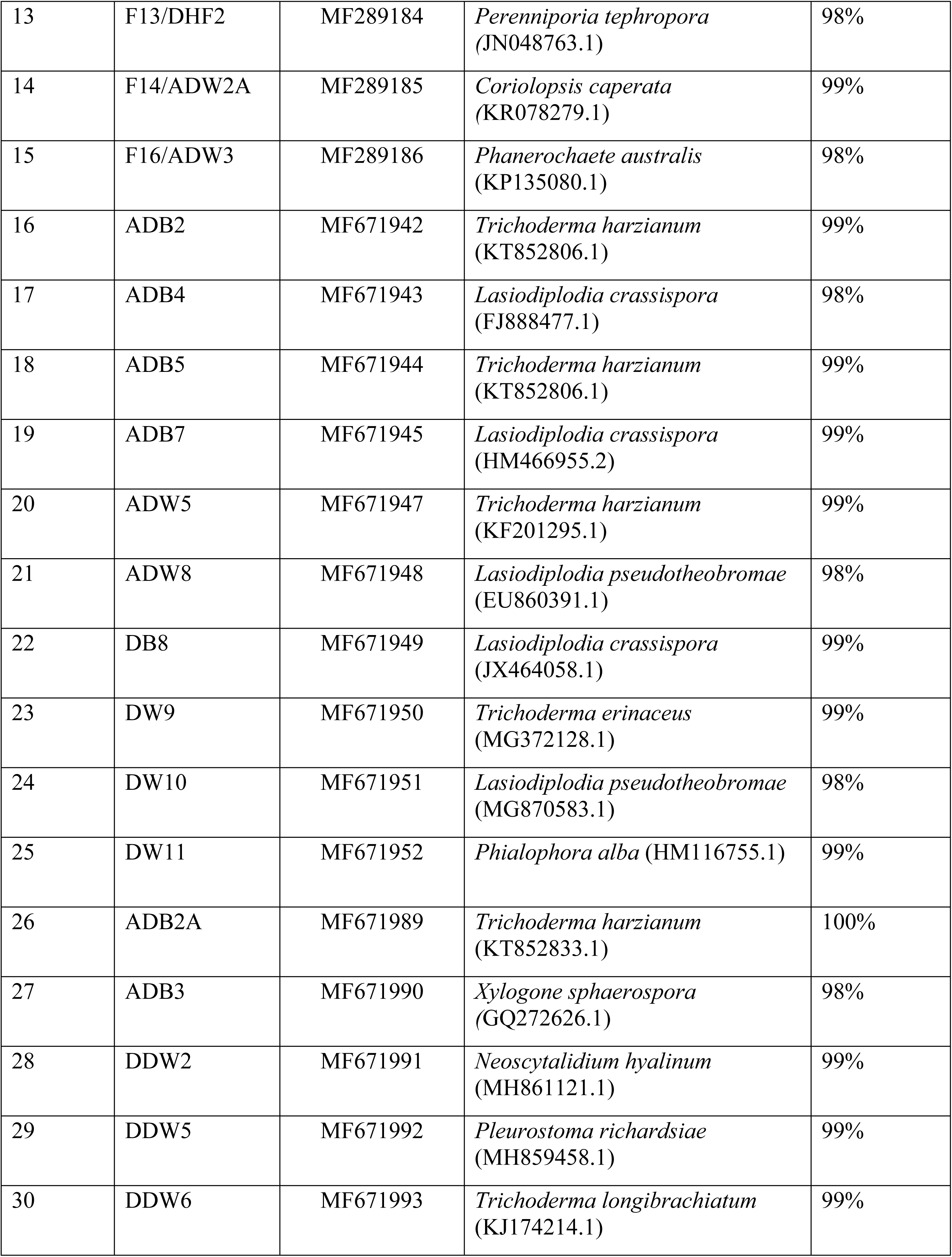
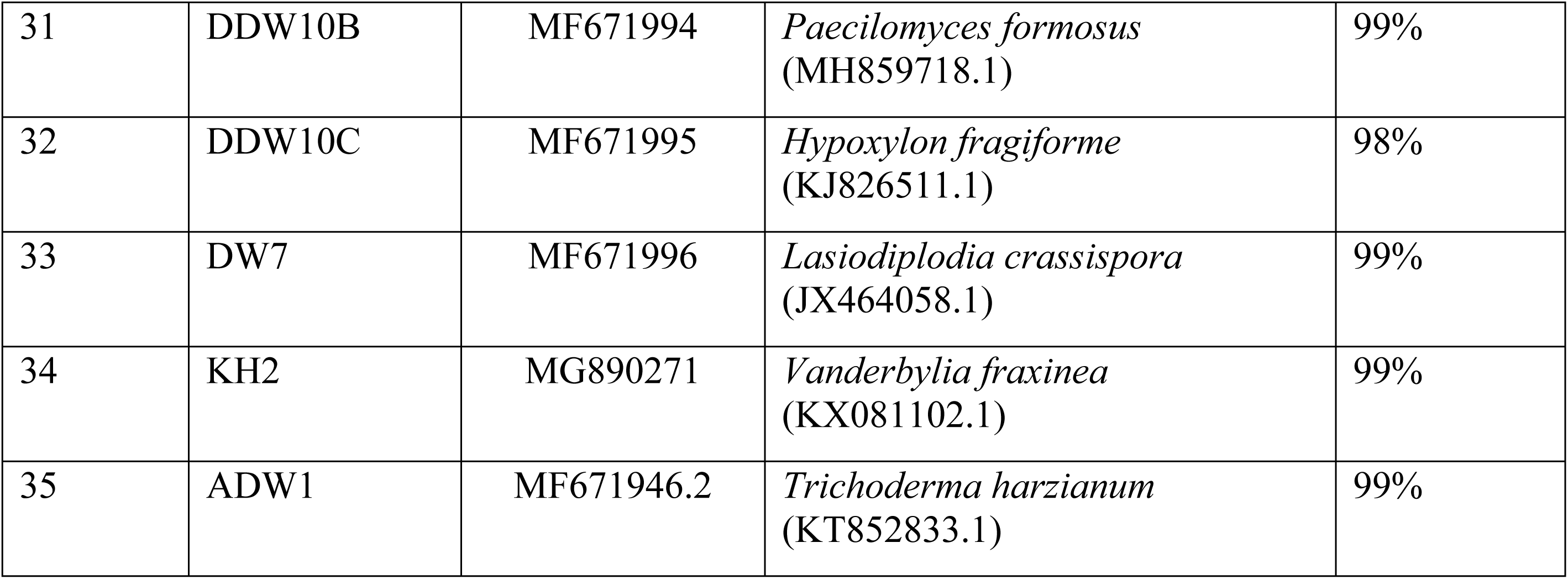
Isolate name, GenBank accession, species name and the GenBank accession number, which showed the highest sequence homology and the percent similarity.

### Molecular phylogenetic analysis

DNA sequences were aligned using a multiple sequence alignment algorithm, Multiple Sequence Comparison by Log-Expectation (MUSCLE) [25], implemented in MEGA ver. X [26]. Gblocks 0.91b was used to eliminate poorly aligned positions and divergent regions of the alignment [27]. All characters were equally weighted and the gaps were treated as missing data. No out group was used and tree was unrooted. The evolutionary history was inferred using several phylogenetic tree construction methods, Maximum Likelihood, maximum parsimony and neighbor-joining, implemented in MEGA ver. X software. For Maximum Likelihood method, initial tree(s) for the heuristic search were obtained automatically by applying Neighbor-Join and BioNJ algorithms to a matrix of pair-wise distances estimated using the Maximum Composite Likelihood (MCL) approach, and then selecting the topology with superior log likelihood value. To test the statistical support of each branch, one thousand boot strap replications were used as implemented in MEGA X.

### Qualitative determination of laccase production by fungal spp

PDA medium containing guaiacol was prepared by adding guaiacol at 0.01% w/v (G5502, Sigma) into PDA before autoclaving. Guaiacol PDA plates were inoculated using a mycelial disk (5 mm diam.) obtained from an actively growing edge of 7 days old culture plates. Triplicates of plates were incubated at room temperature (30-32 °C) for 5 days and the presence of brick red color around the mycelia or underside of the plate was observed [20]. Experiment was repeated once. Isolates that showed consistent red coloration were used for quantitative analysis.

### Quantitative determination of laccase production by fungal spp

Lignocellulose degrading enzyme production was quantitatively determined following [20] for selected 15 samples based on the qualitative assay. Liquid media consisted of 2 % (w/v) wood dust of *Mangifera indica*, glucose, salt solution and Hutner’s trace elements as described in Swe [20]. Erlenmeyer flasks filled with 150.00 mL of liquid media (with saw dust) were autoclaved at 121°C for 20 min. and inoculated with 4 mycelial discs (5.00 mm diam.) obtained from the actively growing edges of 7-day old pure cultures of each fungal isolate. Experiment was conducted in triplicate for the selected 15 fungal isolates and one isolate continuously produced negative results was used as the negative control (Isolate IMF5). In addition, media with plain PDA plugs was also served as a negative control. Flasks were incubated at room temperature (30-32 °C) in a shaker (Stuart-SSL1, UK) (120 rpm) and 1.00 mL of each sample was collected once in the 5^th^, 8^th^ and 12^th^ day during the incubation period. The collected samples were centrifuged at 5100 rpm for 15 min and the supernatant (here after referred as enzyme extract) was stored at −20°C [20]. Reaction mixture (5 mL) contained 3.90 mL acetate buffer (10 mM, pH 5.0), 1.00 mL guaiacol (1.76 mM) and 0.1 mL of the enzyme extract, and incubated at 25°C for 2 hrs. followed by the measuring of absorbance at 450 nm (Thermo scientific Multiskan GO, Finland). In the blank, enzyme extract was substituted with the buffer. Enzyme activity is considered to be corresponding with the increase in absorbance at 450 nm as indicated in Arora and Sandhu [28]. One-way Analysis of Variance (ANOVA) was carried out using Minitab 17 software (Minitab Inc., USA) to determine if there is a significant difference among isolates for the laccase production. Experiment was repeated once.

### Triphenylmethane dye decolorization capacity of laccase producing fungi

Conical flasks containing 25 mL of sterilized medium containing glucose (10 g/L), KH_2_PO_4_ (1 g/L), MgSO_4_ (0.5 g/L), CaCl_2_ (0.14 g/L), yeast extract (1 g/L), thiamine (0.0025 g/L) and bromophenol blue (0.05%) were inoculated with 3 mycelial discs (5.00 mm) of 7-day old actively growing colonies [29]. Flasks were incubated at room temperature for 21 days in the shaker (Stuart-SSL1, UK) at 210 rpm. Two milliliters from each flask was obtained in 8^th^ and 21^st^ days after incubation and centrifuged at 12,000 rpm for 5 minutes to obtain cell free supernatant. Absorbance of each sample was measured at 590 nm (Thermo Scientific Orion aquamate 8000 UV-VIS spectrophotometer, USA) following Nidadavolu *et al.* [30]. Previously used IMF5 was used as the negative control. In addition, media with plain PDA plugs was also served as a negative control. The decolorization efficiency was determined using the following equation [31],

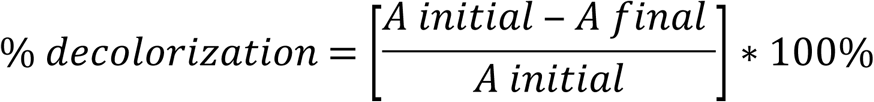

where, *A initial* is the absorbance at the beginning and *A final* is the final absorbance.

To determine the correlation between laccase production and the dye decolonization abilities, Pearson’s correlation was carried out.

## Results

### Isolation of fungi from decaying woods

When PDA was used as the medium, very often fast growing genera of *Trichoderma* and *Lasiodiplodia* spp. were isolated and therefore, semi selective media amended with both antibiotic and benzimidazole was used to capture other genera as well. All together 55 isolates were obtained and most of the colonies that showed *Trichoderma* like morphology were excluded from the subsequent studies since it is well represented in many scientific literatures.

### Ability of isolated fungi to utilize wood as the sole C source

None of the fungal isolates showed significant difference (p<0.05) in their colony diameters on glucose amended vs. glucose non amended media. In some instances, though not significant glucose increased the colony diameters (Fig 2). All the tested isolates were able to utilize wood as their carbon source and based on the colony diameters after three days of incubation, 43 isolates were categorized into three categories as fast growers (>6.0 cm diameter), intermediate growers (6.0-3.0 cm) and slow growers (<3.0 cm). Figure 2 shows the variation in colony diameters on both media for the 43 isolates. Raw data were deposited in Dryad repository https://doi.org/10.5061/dryad.g7f4m0j

**Fig 2.**
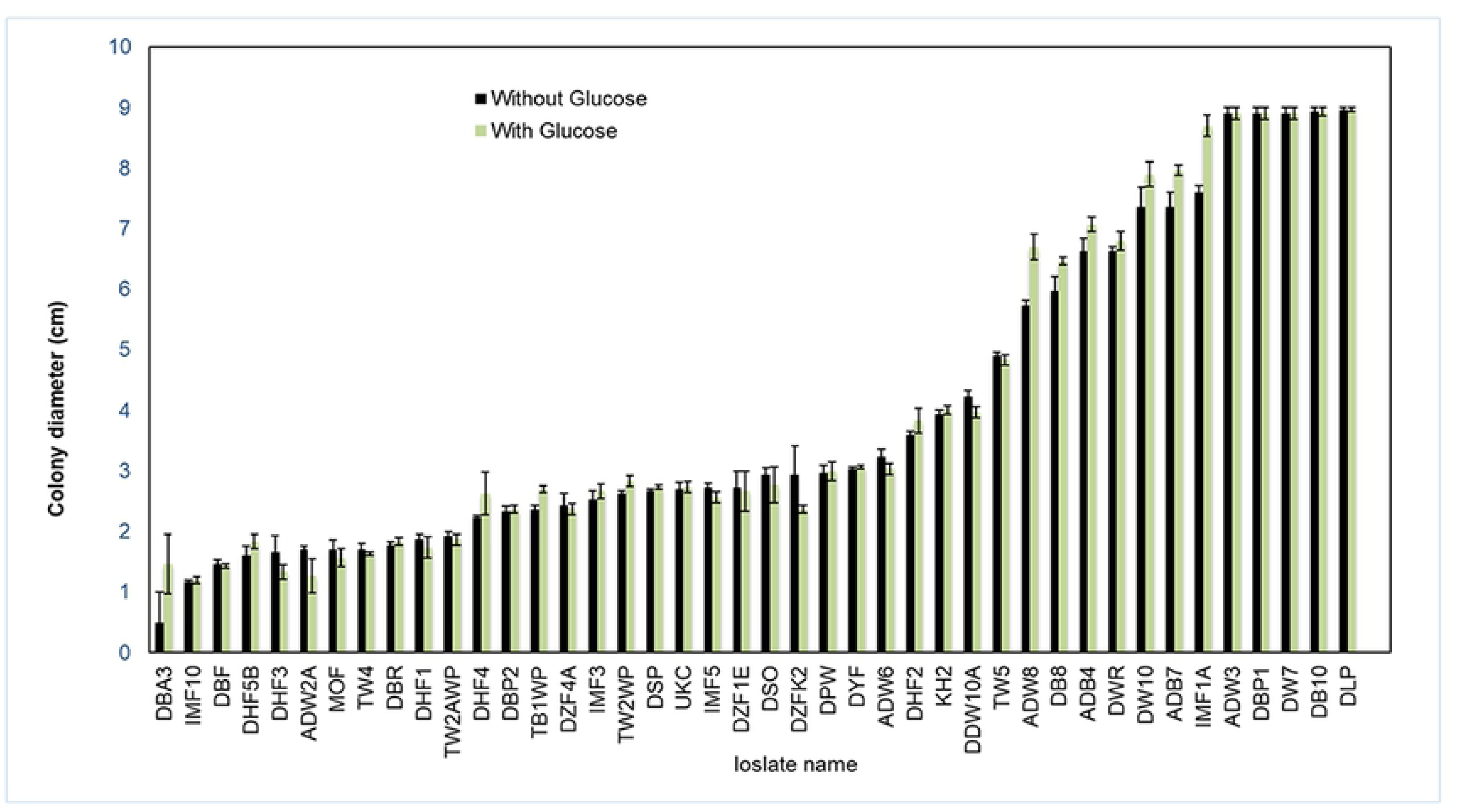
Bar chart showing mean colony diameter (cm) in wood agar (with and without glucose) and names of fungal isolates. Whiskers indicate one SE.

### Identification of fungi

Since most of the fungal isolates were sterile (Fig 3) it was intended to use molecular identification method.

**Fig 3.**
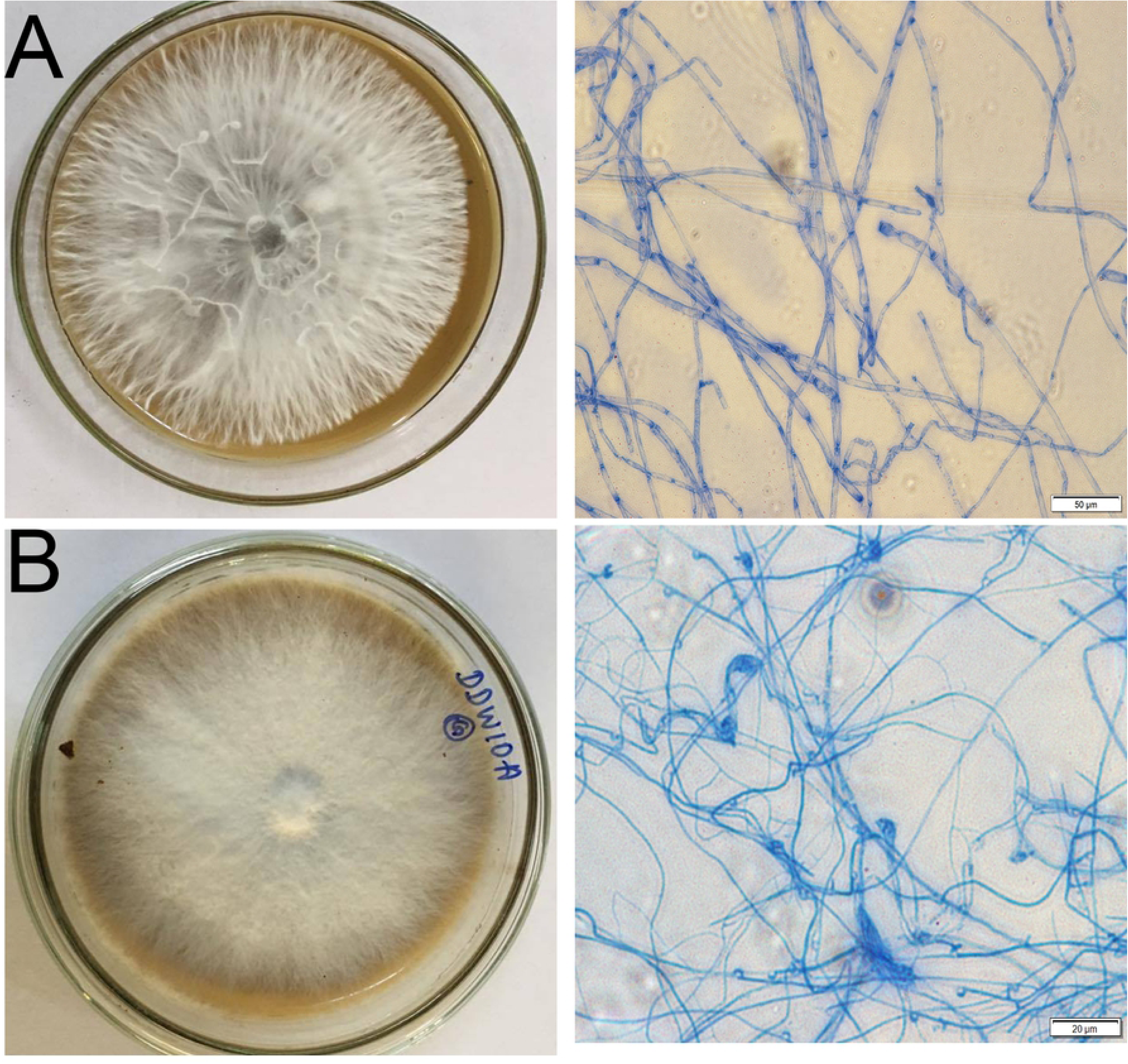
Colony morphology (10-day old culture in the left) and microscopic features of the isolates in the right of the isolates (A) KH2 and (B) DDW10A. Scale bar shows 50 µm and 20 µm respectively.

**Fig 4.**
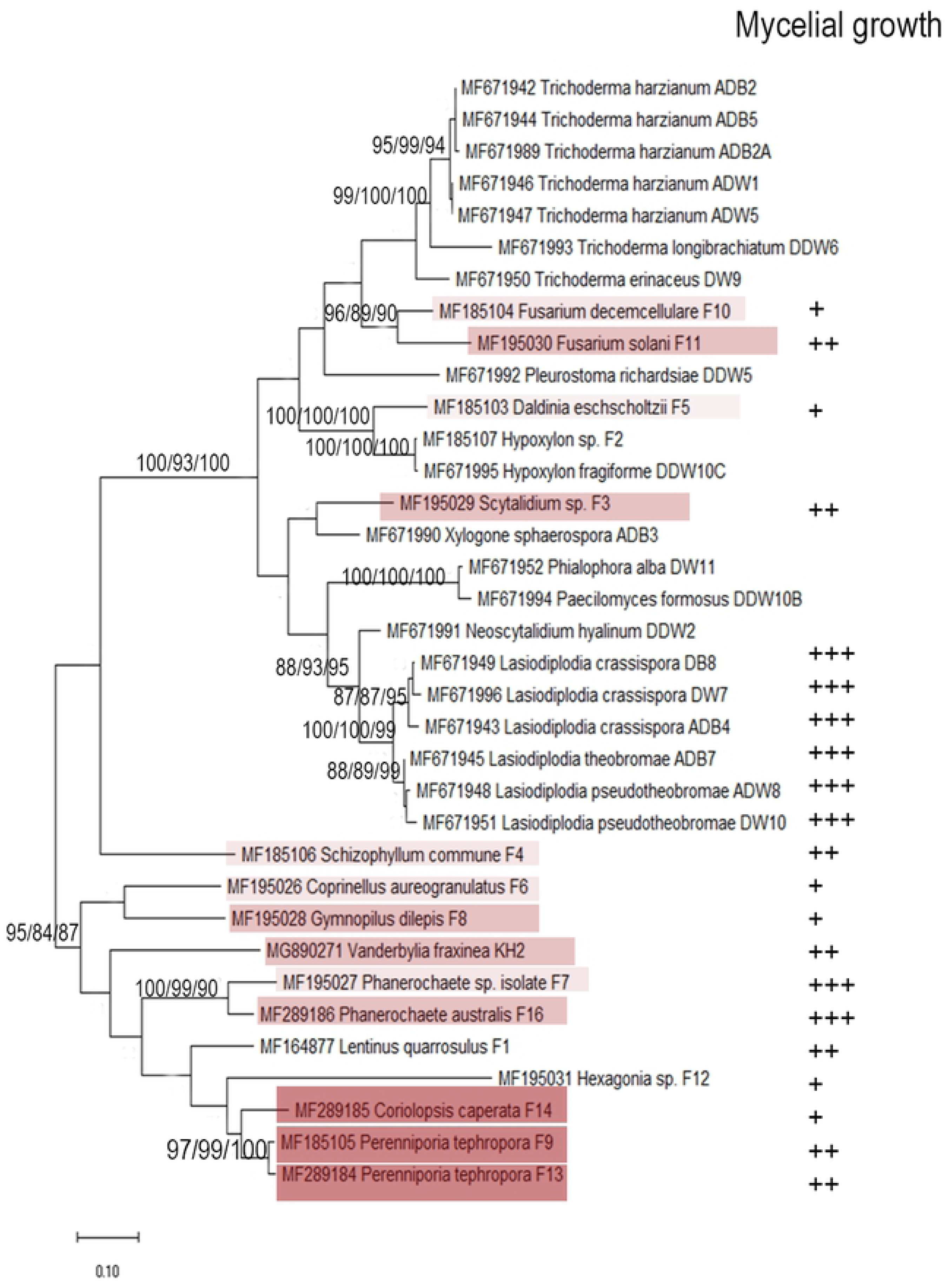
Maximum likelihood tree (unrooted) with the highest log likelihood showing phylogenetic placement of 35 wood decay fungi isolated from Sri Lanka. Bootstrap support is shown above each branch in three different analyses, Maximum Likelihood, Maximum Parsimony and Neighbor Joining respectively. Amount of laccase production in quantitative assay is depicted in red colored boxes in each taxon (darker the color higher the laccase production). Colony diameter is indicated in +, ++ and +++ to represent slow medium and fast growers respectively. Growth diameter was not determined for the isolates with empty cells.

DNA extraction was successful for all the fungal isolates. When ITS1/4 or ITS4/5 primer combinations were used, most of the isolates produced single clear bands. Table 1 shows BLASTn results of each isolate along with the published/vouchered species that showed the highest sequence similarity. After removing most of the fungal cultures with morphologically similar characteristics on PDA, 35 were selected for molecular identification to avoid repetition.

### Molecular phylogenetic analysis

The evolutionary history was inferred by using the Maximum Likelihood method and Tamura-Nei model [26]. Unrooted the tree with the highest log likelihood (−9113.58) is shown in the Fig 3. Initial tree(s) for the heuristic search were obtained automatically by applying Neighbor-Join and BioNJ algorithms to a matrix of pairwise distances estimated using the Maximum Composite Likelihood (MCL) approach, and then selecting the topology with superior log likelihood value. In 35 isolates, there were a total of 654 positions in the final dataset. The percentage of trees in which the associated taxa clustered together is shown next to the branches for maximum likelihood, Maximum parsimony and neighbor joining methods respectively. Colony diameter data were also included and represented by +, ++ or +++ to indicate slow medium and fast growth respectively.

### Qualitative determination of laccase production by fungal spp

Out of 43, only 15 isolates were observed to be consistently producing laccase as indicated by the production of dark red/brown color development in the PDA medium amended with guaiacol in several repeated experiments. Evaluated fungal strains produced varying levels of color intensities as well as coloration patterns while negative control remained without any red color development.

### Quantitative determination of laccase production by fungal spp

Isolates were significantly different from each other in their mean absorbance values as measured in 5^th^, 8^th^ and 12^th^ days indicating significant difference in laccase production (p<0.05) (Table 2). Both of the isolates of *Perenniporia tephropora*, and the rest of the isolates of *Coriolopsis caperata, Gymnopilus dilepis, Fusarium solani, Vanderbylia fraxinea* and *Scytalidium* sp. were among the top seven laccase producers in both trials.

**Table 2:**
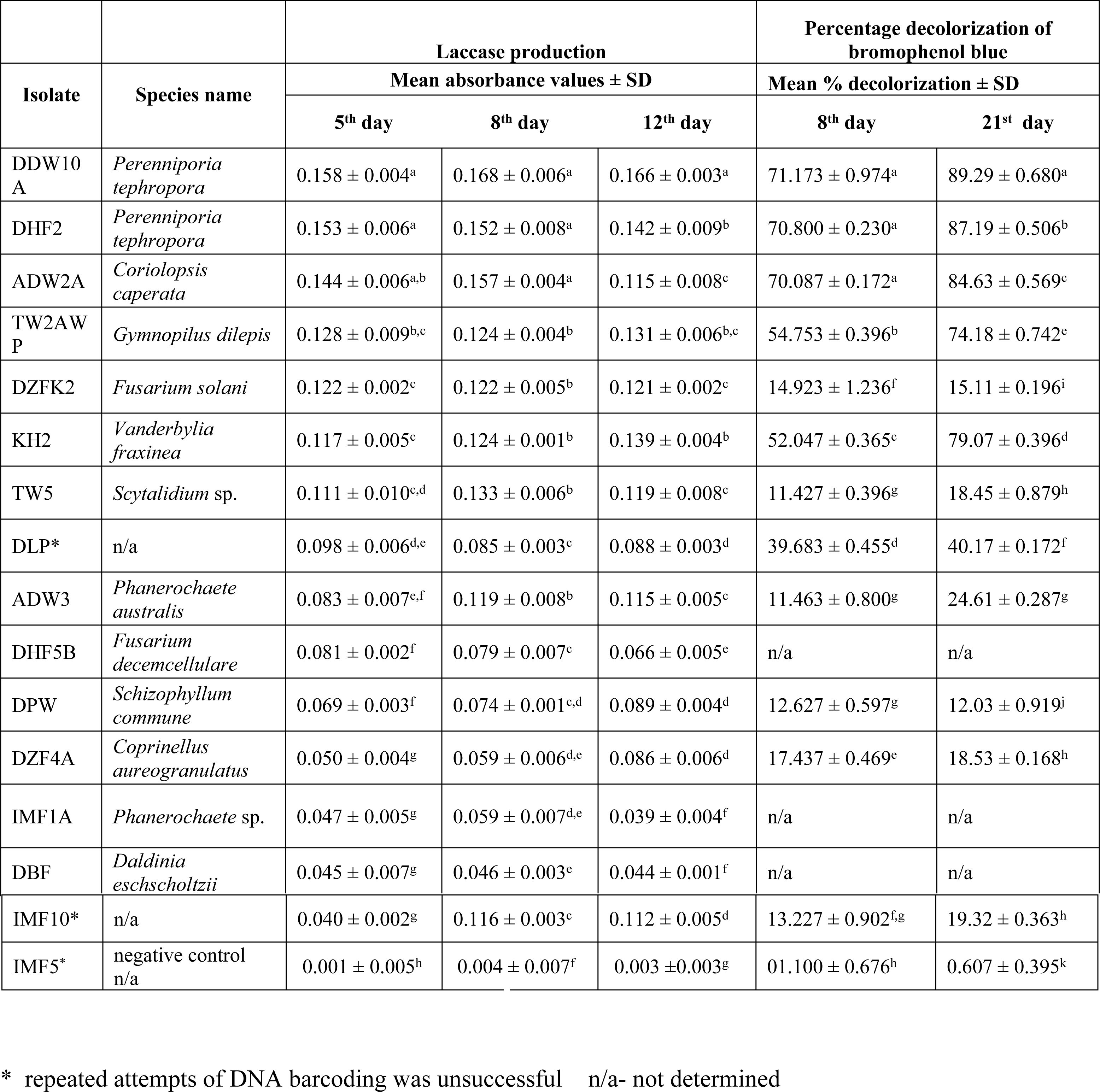
Varying amounts of laccase production as indicated by the absorbance values and percentage dye decolorization (± SD) of the isolates. Isolates that do not share the same letter were significantly different from each other (p<0.05).

### Triphenylmethane dye decolorization assay

Fifteen laccase producers were selected for the determination of their ability to decolorize one of the triphenylmethane dye, bromophenol blue and the results of the quantitative were shown in the Table 2. Isolates were significantly different for the dye decolorization ability as measured by the absorbance values (p<0.05). While both *Perenniporia tephropora* species and *Coriolopsis caperata* isolate decolorized more than 70% of the dye by the 8^th^ day, *Vanderbylia fraxinea* and *Gymnopilus dilepis* isolates decolorized the dye by 70% after three weeks. Each treatment in triplicates showed the decolorization of the dye and no decolorization was observed in negative controls (Table 2). Qualitative test results were indicative of quantitative results. Other laccase positive fungal isolates displayed low decolorization efficiency (below 50%). Non-laccase producer, IMF5, had the least decolorization efficiency (below 10%). According to the Pearson’s correlation analysis, there was a strong and significant correlation (R = 0.64, P = 0.023) between the dye decolorization and laccase production abilities as measured in absorbance values.

## Discussion

Here we reported the presence of 23 different fungal species present in total of 35 decaying hardwood samples (each with 1 cm^2^) and out of them, 11 species; *Coprinellus aureogranulatus, Coriolopsis caperata, Hypoxylon fragiforme, Neoscytalidium hyalinum, Perenniporia tephropora, Phanerochaete australis, Phialophora alba, Pleurostoma richardsiae, Paecilomyces formosus, Vanderbylia fraxinea* and *Xylogone sphaerospora*, were the first reports of Sri Lanka. This finding is important since it highlights that Sri Lankan fungal diversity studies are lagged behind though it is one of the 34 bio diversity hot spots in the world. Except a single study conducted by Ediriweera *et al.* [32]where 14 out of 49 fungal fruiting bodies were identified up to the species level from ‘Sigiriya’ forest in the dry zone, our study is the first to use molecular level fungal species determination on decaying hardwoods in Sri Lanka. It is also important to note that two isolates, F3/TW5 and F12/DHF1 did not show sufficient sequence homology for any of the published records in the GenBank and tentatively identified as *Scytalidium* sp. (86%) and *Hexaginia* sp. (81%) and warrants further studies.

It was hypothesized that fungal species capable of degrading hardwoods are also high laccase produces. Presence of laccase has been determined by the formation of reddish brown color due to oxidation of guaiacol by laccase in many studies [33,34] and was implemented in the present study. Among the tested species, *Perenniporia tephropora* produced the highest amount of laccase and this finding is in line with Younes *et al.* [35]. There was no significant difference between *P. tephropora* and *Coriolopsis caperata* for the laccase production. Other best laccase producers were *Gymnopilus dilepis, Fusarium solani* and *Vanderbylia fraxinea.* Though the mother plant species of most of the decayed wood pieces were not known, it is interesting to note that *P. tephropora* and *V. fraxinea* were isolated from decaying woods of two economically and medicinally important hardwood species, Ebony (*Diospyros ebenum*) and Neem (*Azadirachta indica*) respectively. *Diospyros ebenum* also known as black wood is native to India and Sri Lanka and belongs to one of the hardest wood species resistant to withering and fungal infections. Neem is also known to contain numerous chemical compounds with medicinal importance and also known as the tree of the 21^st^ century by the United Nations. Specially we tested whether these two fungi can utilize *Diospyros ebenum* wood dust as the sole C source and confirmed to be able to grow on wood agar palates (observational study only). Fungi isolated from the decaying woods of these plants were among the top laccase producers supporting our original hypothesis. Since Hori et al. (2017) found that white rot polyporales show greater enzymatic diversity than brown rot polyporales [36] and both these fungal species were isolated from white rot hardwoods, it is worth conducting a full genomic analysis.

We found that the laccase production was positively correlated with triphenylmethane dye decolorization. Laccase and other ligninase enzyme extract’s direct involvement in dye decolorization has been previously reported [37,38,39]. We also found that *P. tephropora* and *C. caperata* were the strongest bromophenol blue decolorizers similar to previous studies [35,40,41]. However, to the best of our knowledge triphenylmethane dye decolorization ability and laccase production of *Vanderbylia fraxinea* and *Gymnopilus dilepis* has never been reported in the fungal kingdom before and we report it for the first time.

These findings were of high importance, since it is known that synthetic dyes used in the textile, paper, leather and cosmetic industries are highly toxic, resistant to light, chemical and microbial degradation. Among many types of dyes, triphenylmethane dyes are resistant to enzymatic decolorization and it is time consuming [42]. Laccases require no H_2_O_2_ for oxidation reactions like other oxidases and that makes laccase an important enzyme in bio remediation. In the phylogenetic analysis, though laccase production is not monophylectic, all the high laccase producers were clustered together in a single clade of basidiomycetes. Though Ascomycetes have laccase producing ability, they produce relatively low amounts. It has also been reported that number of laccase gene copies could vary among species depending on the life history traits [43].

Finally, this study reported 11 wood decay fungal species in Sri Lanka as first reports highlighting the importance of a thorough fungal diversity study in Sri Lanka since there is a high potential to report novel biotechnologically important species. It was also proved that most of the decaying wood associated fungi are indeed can utilize wood as the sole C source. To the best of our knowledge, laccase production and triphenylmethane dye decolorization ability of *Vanderbylia fraxinea* and *Gymnopilus dilepis* have never been reported in the fungal kingdom before. It was also found that laccase production was positively correlated with the dye decolozination ability. Therefore, the selected isolates have very high potential in applying for a greener biotechnology.

## Acknowledgements

This research was funded by TWAS and ICGEB research grants.

## References

1. Fox EM, Howlett BJ. Secondary metabolism: regulation and role in fungal biology. Curr Opin Microbiol. 2008 Dec;11(6):481–7.

2. Hawksworth DL. The fungal dimension of biodiversity: magnitude, significance, and conservation. Mycol Res [Internet]. 1991;95(6):641–55. Available from: http://www.sciencedirect.com/science/article/pii/S0953756209808101

3. Blackwell M. The fungi: 1, 2, 3 5.1 million species? Am J Bot. 2011 Mar;98(3):426–38.

4. Aime MC, Brearley FQ. Tropical fungal diversity: closing the gap between species estimates and species discovery. Biodivers Conserv [Internet]. 2012 Aug;21(9):2177–80. Available from: https://doi.org/10.1007/s10531-012-0338-7

5. Hawksworth D. The tropical fungal biota: census, pertinence, prophylaxis, and prognosis. Aspects of tropical mycology. Symposium, Liverpool, 1992. 1993. 265–293 p.

6. Gunatilleke Nimal, Pethiyagoda Rohan G. Special issue inside.pdf. 2008;(25):25–61.

7. Wilkin P, Dassanayake MD, Clayton WD. A Revised Handbook to the Flora of Ceylon. Volume XIV. Kew Bull. 2007 Nov 27;55(4):1015.

8. Karunarathna S. Current status of knowledge of Sri Lankan mycota. Curr Res Environ Appl Mycol. 2018;2(1):18–29.

9. A Blanchette R. Delignification by Wood-Decay Fungi. Vol. 29, Annual Review of Phytopathology. 2003. 381–403 p.

10. Margot J, Bennati-Granier C, Maillard J, Blanquez P, Barry DA, Holliger C. Bacterial versus fungal laccase: potential for micropollutant degradation. AMB Express. 2013 Oct;3(1):63.

11. Jayasuriya AHM, Kitchener DJ, Biradar CM. Viability status of biosphere reserves in Sri Lanka. J Natl Sci Found Sri Lanka. 2011;39(4):303–19.

12. Iqbal MCM, Nalaka GDA, Kumarathunage MDP. Tree diversity in a tropical dry mixed evergreen forest plot in Sri Lanka. Proc Int For Environ Symp. 2012;17(August 2015):2012.

13. Madurapperuma BD, Oduor PG, Kuruppuarachchi KAJM, Wijayawardene DNN, Munasinghe JU. Comparing Floristic Diversity between a Silviculturally Managed Arboretum and a Forest Reserve in Dambulla, Sri Lanka. J Trop For Environ. 2018;3(2):11–22.

14. Upadhyay P, Shrivastava R, Agrawal PK. Bioprospecting and biotechnological applications of fungal laccase. 3 Biotech. 2016;6(1):1–12.

15. Jeon J-R, Baldrian P, Murugesan K, Chang Y-S. Laccase-catalysed oxidations of naturally occurring phenols: from in vivo biosynthetic pathways to green synthetic applications. Microb Biotechnol. 2012 May;5(3):318–32.

16. Rodriguez E, Pickard MA, Vazquez-Duhalt R. Industrial dye decolorization by laccases from ligninolytic fungi. Curr Microbiol. 1999 Jan;38(1):27–32.

17. Liu H, Cheng Y, Du B, Tong C, Liang S, Han S, et al. Overexpression of a novel thermostable and chloride-tolerant laccase from Thermus thermophilus SG0.5JP17-16 in Pichia pastoris and its application in synthetic dye decolorization. PLoS One. 2015;10(3):e0119833.

18. Surwase S V, Patil SA, Srinivas S, Jadhav JP. Interaction of small molecules with fungal laccase: A Surface Plasmon Resonance based study. Enzyme Microb Technol. 2016 Jan;82:110–4.

19. Baumgartner K, Travadon R, Bruhn J, Bergemann SE. Contrasting patterns of genetic diversity and population structure of Armillaria mellea sensu stricto in the eastern and western United States. Phytopathology. 2010 Jul;100(7):708–18.

20. Swe KT. Screening of potential lignin-degrading microorganisms and evaluating their optimal enzyme producing culture conditions. 2011;86.

21. Cenis JL. Rapid extraction of fungal DNA for PCR amplification. Nucleic Acids Res. 1992;20(9):2380.

22. White TJ, Bruns T, Lee S, Taylor J. AMPLIFICATION AND DIRECT SEQUENCING OF FUNGAL RIBOSOMAL RNA GENES FOR PHYLOGENETICS. PCR Protoc [Internet]. 1990 Jan 1 [cited 2019 May 18];315–22. Available from: https://www.sciencedirect.com/science/article/pii/B9780123721808500421?via%3Dihub

23. Maduranga K, Attanayake RN, Santhirasegaram S, Weerakoon G, Paranagama PA. Molecular phylogeny and bioprospecting of Endolichenic Fungi (ELF) inhabiting in the lichens collected from a mangrove ecosystem in Sri Lanka. PLoS One [Internet]. 2018 Aug 29;13(8):e0200711. Available from: https://doi.org/10.1371/journal.pone.0200711

24. Hall T. BioEdit: A User-Friendly Biological Sequence Alignment Editor and Analysis Program for Windows 95/98/NT. Vol. 41, Nucleic Acids Symposium Series. 1999. 95–98 p.

25. Edgar RC. MUSCLE: multiple sequence alignment with high accuracy and high throughput. Nucleic Acids Res. 2004;32(5):1792–7.

26. Kumar S, Stecher G, Li M, Knyaz C, Tamura K. MEGA X: Molecular Evolutionary Genetics Analysis across Computing Platforms. Mol Biol Evol. 2018 Jun;35(6):1547–9.

27. Talavera G, Castresana J. Improvement of phylogenies after removing divergent and ambiguously aligned blocks from protein sequence alignments. Syst Biol. 2007 Aug;56(4):564–77.

28. Arora DS, Sandhu DK. Decomposition of angiospermic wood sawdust and laccase production by two Pleurotus species. J Basic Microbiol [Internet]. 1987 Jan 1;27(4):179–84. Available from: https://doi.org/10.1002/jobm.3620270402

29. Jonathan SG, Fasidi IO. Effect of carbon, nitrogen and mineral sources on growth of Psathyerella atroumbonata (Pegler), a Nigerian edible mushroom. Food Chem [Internet]. 2001;72(4):479–83. Available from: http://www.sciencedirect.com/science/article/pii/S030881460000265X

30. Nidadavolu SVSSSLHB, Gudikandula K, Pabba SK, Maringanti SC. Decolorization of triphenyl methane dyes by *Fomitopsis feei* Nat Sci. 2013;05(06):30–5.

31. Lopez C, Moreira MT, Feijoo G, Lema JM. Dye decolorization by manganese peroxidase in an enzymatic membrane bioreactor. Biotechnol Prog. 2004;20(1):74–81.

32. Ediriweera SS, Wijesundera RLC, Weerasena OVDSJ. Macrofungi from the Sigiriya wilderness in Sri Lanka. 2014;52(1):47–51.

33. Nishida, T.; Kashino, Y.; Mimura, A.; Takahara Y. Lignin biodegradation by wood-rotting fungi I. Screening of lignin-degrading fungi. Mokuzai Gakkaishi J Japan Wood Res Soc [Internet]. 1988 Jan 1;34(6):530–6. Available from: https://eurekamag.com/research/001/628/001628736.php

34. Luterek J, Gianfreda L, Wojtaś-Wasilewska M, Rogalski J, Jaszek M, Malarczyk E, et al. Screening of the wood-rotting fungi for laccase production: Induction by ferulic acid, partial purification, and immobilization of laccase from the high laccase-producing strain, Cerrena unicolor. Vol. 46, Acta Microbiologica Polonica. 1997. 297–311 p.

35. Ben Younes S, Mechichi T, Sayadi S. Purification and characterization of the laccase secreted by the white rot fungus Perenniporia tephropora and its role in the decolourization of synthetic dyes. J Appl Microbiol. 2007 Apr;102(4):1033–42.

36. Hori C, Gaskell J, Igarashi K, Samejima M, Hibbett D, Henrissat B, et al. Genomewide analysis of polysaccharides degrading enzymes in 11 white- and brown-rot Polyporales provides insight into mechanisms of wood decay. Mycologia [Internet]. 2013 Nov 1;105(6):1412–27. Available from: https://doi.org/10.3852/13-072

37. Abd El Monssef RA, Hassan EA, Ramadan EM. Production of laccase enzyme for their potential application to decolorize fungal pigments on aging paper and parchment. Ann Agric Sci [Internet]. 2016;61(1):145–54. Available from: http://www.sciencedirect.com/science/article/pii/S057017831500055X

38. Ratanapongleka K, Phetsom J. Decolorization of Synthetic Dyes by Crude Laccase from Lentinus Polychrous Lev. Vol. 5, International Journal of Chemical Engineering and Applications. 2014. 26–30 p.

39. Qin X, Zhang J, Zhang X, Yang Y. Induction, purification and characterization of a novel manganese peroxidase from Irpex lacteus CD2 and its application in the decolorization of different types of dye. PLoS One. 2014;9(11):e113282.

40. S T, V P, Ingale S, A G. UTILIZATION OF LIGNOCELLULOSES FOR THE PRODUCTION OF LIGNINOLYTIC ENZYMES BY WHITE ROT BASIDIOMYCETE CORIOLOPSIS CAPERATA AGST2. Vol. 2, International Journal of Biotechnology and Biosciences. 2012. 228–238 p.

41. Nandal P, Ravella SR, Kuhad RC. Laccase production by Coriolopsis caperata RCK2011: optimization under solid state fermentation by Taguchi DOE methodology. Sci Rep. 2013;3:1386.

42. Forootanfar H, Moezzi A, Aghaie-Khozani M, Mahmoudjanlou Y, Ameri A, Niknejad F, et al. Synthetic dye decolorization by three sources of fungal laccase. Iranian J Environ Health Sci Eng [Internet]. 2012 Dec 15;9(1):27. Available from: https://www.ncbi.nlm.nih.gov/pubmed/23369690

43. Cázares-García SV, Vázquez-Garcidueñas S, Vázquez-Marrufo G. Structural and phylogenetic analysis of laccases from Trichoderma: a bioinformatic approach. PLoS One [Internet]. 2013 Jan 31;8(1):e55295.#x2013;e55295. Available from: https://www.ncbi.nlm.nih.gov/pubmed/23383142

